# Rat Anterior Insula Symmetrically Represents Tickling-Induced Playful Emotions

**DOI:** 10.1101/2024.04.02.587725

**Authors:** Sarah Dagher, Shimpei Ishiyama

## Abstract

Social play, an integral aspect of animal behavior, is inherently associated with positive emotions, yet the neuronal underpinnings of these playful states remain inadequately explored. We examined the anterior insula’s involvement in processing tickle-induced playful emotions in rats. Our findings revealed diverse patterns of insular activity during tickling, with 20% of the recorded units displaying strong activation, and another 20% exhibiting inhibition. These units responded similarly to other playful contexts, such as gentle touch and hand chase, but not to neutral locomotion. Tickle-activated units demonstrated a positive correlation of firing rates with appetitive vocalization rates, whereas tickle-inhibited units showed a negative correlation. Distinct spike waveforms were associated with the tickle response patterns, suggesting potential cell-type dependencies. However, pharmacological manipulation of the global anterior insula did not yield observable effects on play behavior in rats. Anterograde tracing revealed extensive insular projections to areas including the amygdala and nucleus accumbens. Taken together, our findings suggest that the anterior insula symmetrically represents tickle-induced playful emotional states.

## Introduction

Social play behavior is a fundamental aspect of animal behavior, serving crucial roles in social development, learning, and synapse formation [1-6]. Numerous studies have highlighted several brain regions important in play behavior, including the habenula, prefrontal cortex, nucleus accumbens, periaqueductal gray, and striatum [7-10]. Across various species, social play has been studied through behavioral analysis in various species including hamsters [11, 12], dogs [13], and cats [14], in addition to extensive neuropharmacological investigations in rats [15-17]. In fact, Jaak Panksepp proposed that PLAY is one of the primary emotional systems in the brain, underscoring its pivotal role in mammalian social and emotional development [18].

Laughter-inducing tickling, or gargalesis [19], is a common aspect of social play in humans. Intriguingly, gargalesis is not exclusive to humans but is also observed in other species, including primates [20], and rats [21]. As a result, tickling has emerged as a compelling paradigm for studying playful behaviors in animals, in particular rats [21-25]. When tickled, rats emit 50 kHz ultrasonic vocalizations (USVs), reflecting appetitive emotional states [26]. Previous research, particularly investigations into the somatosensory cortex (S1) [27, 28], the prefrontal cortex [29] and the periaqueductal gray (PAG) [10], has provided insights into the neural substrates underlying tickle-induced behaviors and USVs, demonstrating the activation of these areas in response to tickling. However, despite this progress, the emotional processing of playfulness has remained largely unexplored.

Emotions have long captivated researchers, with numerous theories and debates on their definition, yet there is a consensus that emotions can be characterized by key features, including generalization, persistence, specificity, and scalability [30]. Researchers have achieved significant advancements in understanding their categorization and the brain regions involved in their expression in particular the limbic system [31-34]. Much of the focus in affective neuroscience has traditionally been on negative emotions such as fear and anxiety [35-40], providing valuable insights into the neural substrates of aversive states. Recently, the interest has expanded to encompass positive emotions. Studies employing well-controlled methods such as sucrose consumption have been foundational in this exploration [41, 42]. Yet, these models might not fully encompass the spontaneous and social nature of behaviors associated with positive emotions observed in social animals. As such, there remains a notable gap in our understanding of how the brain processes positive emotions, particularly in the context of naturalistic behaviors like play. Studying how animals express emotions during spontaneous, untrained behaviors like play, not only enhances our understanding of animal cognition and social interaction but also provides valuable parallels to human emotional experiences.

A prominent area that received much attention in the study of emotions is the insular cortex. The insula serves as a central hub for emotional processing, intricately connected with somatosensory cortices [43]. Notably, it projects extensively to regions implicated in reward and motivation, including the amygdala, nucleus accumbens, and ventral tegmental area. Furthermore, observations from patients with lesions in the anterior insula demonstrate pain asymbolia, where they recognize painful sensations but do not perceive them as aversive [44], suggesting the role of the insula in associating tactile sensations with emotions. Conversely, studies have also demonstrated that pleasant touch on the skin can activate the insula [45], highlighting its involvement in processing positive stimuli. Additionally, stimulation of the anterior insula has been shown to elicit laughter in humans [46], further emphasizing its role in emotional expression. However, cellular mechanisms of naturalistic positive emotions such as playfulness within the insula remain elusive. We hypothesise that the anterior insula serves as a neural substrate for representing tickle-induced positive and playful emotions, demonstrating features consistent with typical emotional responses.

Our results show a wide array of physiological responses and neuronal diversity within the anterior insular cortex. Going deeper into specific insular circuits, we highlighted potential involvement of diverse brain regions, including the nucleus accumbens, thalamus, and amygdala, in possibly mediating the emotional responses during social play.

## Materials and Methods

### Animals

All experimental protocols adhered to German regulations on animal welfare (Permit number G 20-1-082). Male juvenile Long-Evans rats, aged 3 weeks upon arrival, were obtained from Janvier. The rats were single housed with unrestricted access to food and water and were kept in a 12-hour light/dark cycle. Experimental procedures, including behavioral habituation, surgeries, and recordings, were conducted during the dark phase.

### Setup and synchronization

The experimental arena consisted of a plastic cylinder measuring 45 cm diameter × 45 cm height used for tickling, with two cameras and microphone attached. The walls were covered with black polyurethane foam. Throughout the sessions, the environment was maintained at a subdued illumination of 20 lx, as measured at the center of the floor. Synchronization of audio, video, and electrophysiological recordings was achieved using TTL pulses, as previously described [47]. The setup is illustrated in Figure 1A.

**Figure 1:**
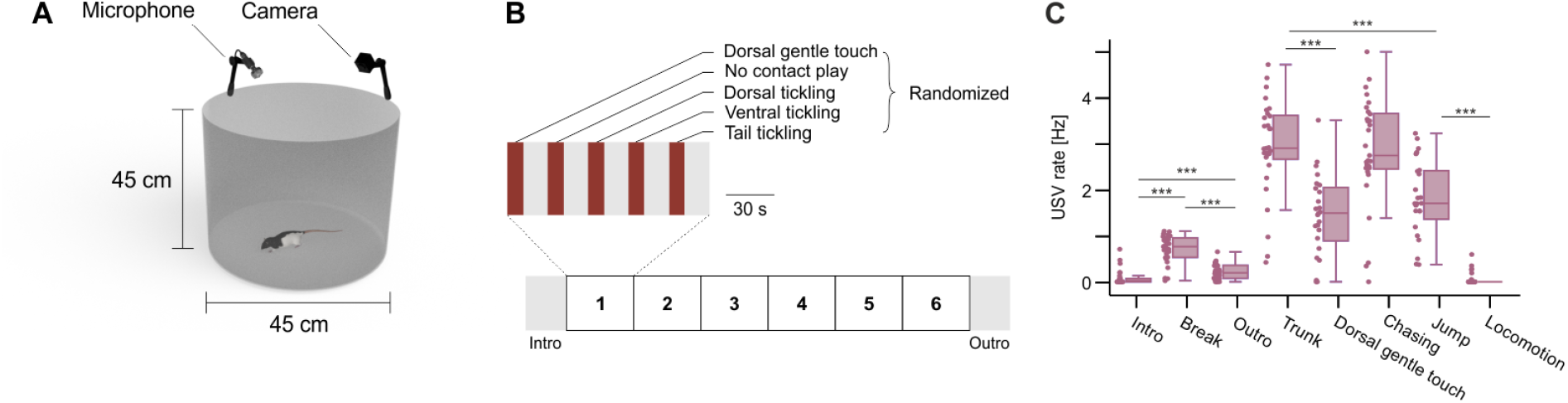
Experimental paradigm and behavioral response. **A)** Tickle arena with simultaneous audio and video recording. **B)** Behavioral paradigm. **C)** Ultrasonic vocalization (USV) rate during different events. Data are obtained from 29 sessions, 7 animals. Statistical comparison was performed using signed-rank test.

### Behavioral paradigm

Preceding electrophysiology and pharmacology experiments, the animals underwent a daily regimen of 10-minute tickling sessions over the course of two weeks to facilitate habituation. Subsequently, tickling experiments were conducted daily for approximately two weeks. The experimental paradigm used for electrophysiology and pharmacology recordings consisted of a 2-min introductory period, followed by six blocks of heterospecific interaction phases. Each block comprised randomized sequences of five distinct phases: dorsal tickling, dorsal gentle touch, ventral tickling, tail tickling, and a no-contact play phase where the hand was introduced into the arena without making physical contact, allowing the animal to engage in chasing behavior. Phases within each block lasted for 15 seconds, interspersed with 10-second interphase break periods. Following the completion of the sixth block, animals were kept in the box for a 2-min concluding period (outro) (Figure 1B). Video and calls were consistently recorded throughout all sessions.

### Sound recording and analysis

Using a condenser ultrasound microphone (CM16/CMPA, Avisoft Bioacoustics, Berlin, Germany) positioned on top of the behavioral box, ultrasonic vocalizations (USVs) were recorded at a sampling rate of 256 kHz and a 16-bit resolution using Avisoft-RECORDER USGH software (Avisoft Bioacoustics, Berlin, Germany). USVs were detected and curated using DeepSqueak [48].

### Video recording and analysis

Video recordings were obtained using two BlackFlyS-U3-28S5C-C cameras (Teledyne FLIR, USA), each operating at 30 frames per second (fps), with one camera positioned above and the other beside the experimental arena. Analysis of the video footage was conducted using BORIS software [49] to annotate the beginnings and the ends of experimental paradigm phases and animal behaviors.

### Electrophysiology experiment

#### Microdrive and implantation

Freely-moving *in vivo* electrophysiological recordings were performed using either tetrodes or silicon probes. For tetrode recordings, we used a Halo-10 microdrive (Neuralynx Inc, USA) consisting of eight independently-movable tetrodes. Tetrodes were made from 12.5 µm diameter Teflon-coated nichrome wire (Sandvik, Sandviken, Sweden) and gold plated to reach an impedance of around 250 kΩ using NanoZ (White Matter LLC, USA). For tetrode tracks identification in the brain, tetrodes were stained with fluorescent dye DiI or DiO (ThermoFisher Scientific Inc., USA) just before implantation. For silicon probe recordings, a E32+R-100-S1-L10 NT probe (ATLAS Neuro, Belgium) consisting of 4 shanks with eight channels per shank mounted on a custom-built microdrive and a head-mount faraday cage, were used in the study. The ground wire of the silicon probe was soldered onto the faraday cage. The shanks were also stained with DiI before implantation. No gold plating was performed for silicon probes.

After two weeks of habituation, animals were anesthetized using isoflurane (4% for induction, 1.5-3% for maintenance; cp-pharma, Burgdorf, Germany), and securely positioned in a stereotaxic apparatus (Stoelting, Wood Dale, IL, USA). Body temperature was maintained with a heating pad and monitored by a rectal probe (Stoelting, Wood Dale, IL, USA). Lidocaine was administered via subcutaneous injection into the scalp, and carprofen (10%, 5 mg/kg body weight; Zoetis, Germany) was subcutaneously injected in the dorsal trunk prior to making an incision. Following skull scraping, a stainless-steel screw and two screws with a silver wire soldered to them were affixed to the skull as an anchor and grounds, respectively. The skull surface was cleaned with 1% hydrogenperoxide and 70% ethanol. The surface was then treated with Optibond (Kerr Italia, Salerno, Italy) and covered with Charisma dental filling (Heraeus Kulzer, Hanau, Germany). After craniotomy above the left anterior insula (2.76 mm anterior, 4.5 mm lateral from Bregma) and durotomy, the microdrive was positioned onto the brain, and the exposed brain area was covered with Dura-Gel (Cambridge NeuroTech, UK). The silver wires attached to the skull screws were connected to the microdrive faraday cage. Securing of the microdrive was fixed using dental cement (Paladur, Heraeus Kulzer, Hanau, Germany). Upon completion of the surgery, carprofen (at the same dosage as previously mentioned) was subcutaneously administered before the animal woke up.

#### Electrophysiological recordings

Starting two days after surgery, tetrodes/silicon probe were lowered into the brain by ∼0.25 mm daily. Upon detection of neuronal spikes, experimental procedures started, ensuring a minimum of 1 hour elapsed after the tetrodes/probe were lowered to allow for stabilization of tissue drift. Extracellular spikes originating from the left anterior insular cortex were recorded at 30 kHz sampling rate and bandpass-filtered between 0.6 and 6 kHz using Cheetah software, and LabLynx (Neuralynx, Bozeman, MT, USA).

#### Spike analysis

Spike sorting and clustering were performed using JRCLUST [50]. Units with average firing rate < 0.1 Hz were excluded. Units that lack temporal stability across the recording (absolute correlation coefficient between time and firing rate > 0.4) were excluded. Peristimulus time histograms (PSTHs) of firing rates were aligned to the offset of events rather than the onset, due to the preceding physical restraint and pinning of the rats prior to ventral tickling, ensuring that the period following the offset represents a true absence of external stimuli. For population analyses, firing rates of each unit were standardized in terms of z-score, based on the mean and standard deviation of firing rate (0.5 s bin) in the period out of the event i.e. 0 till end of PSTH time range. Accordingly, units with no spike during this period were excluded from the analysis. Population PSTHs were smoothed with Gaussian filter and represented as mean ± SEM.

Hierarchical clustering of PSTHs was performed to categorize units exhibiting similar firing patterns. The Euclidean distance was employed to measure cluster distances and Ward algorithm was utilized to guide the agglomerative clustering process. A threshold to cut the hierarchical tree was defined at the longest gap in the dendrogram.

To assess the relationship between USVs and firing rates, a linear mixed-effects model was employed, treating the unit identifier as a random intercept.

For waveform analysis, mean spike waveform of each unit was oversampled at 5x using spline-based interpolation [51]. The oversampled waveforms were then normalized to their peaks, and waveform features including peak-to-trough ratio, peak-to-trough time, repolarization time, half-maximum width, and the area under curve (AUC) were calculated for each unit (Figure 2J) [52] before being projected into UMAP space and clustered using K-Means.

**Figure 2:**
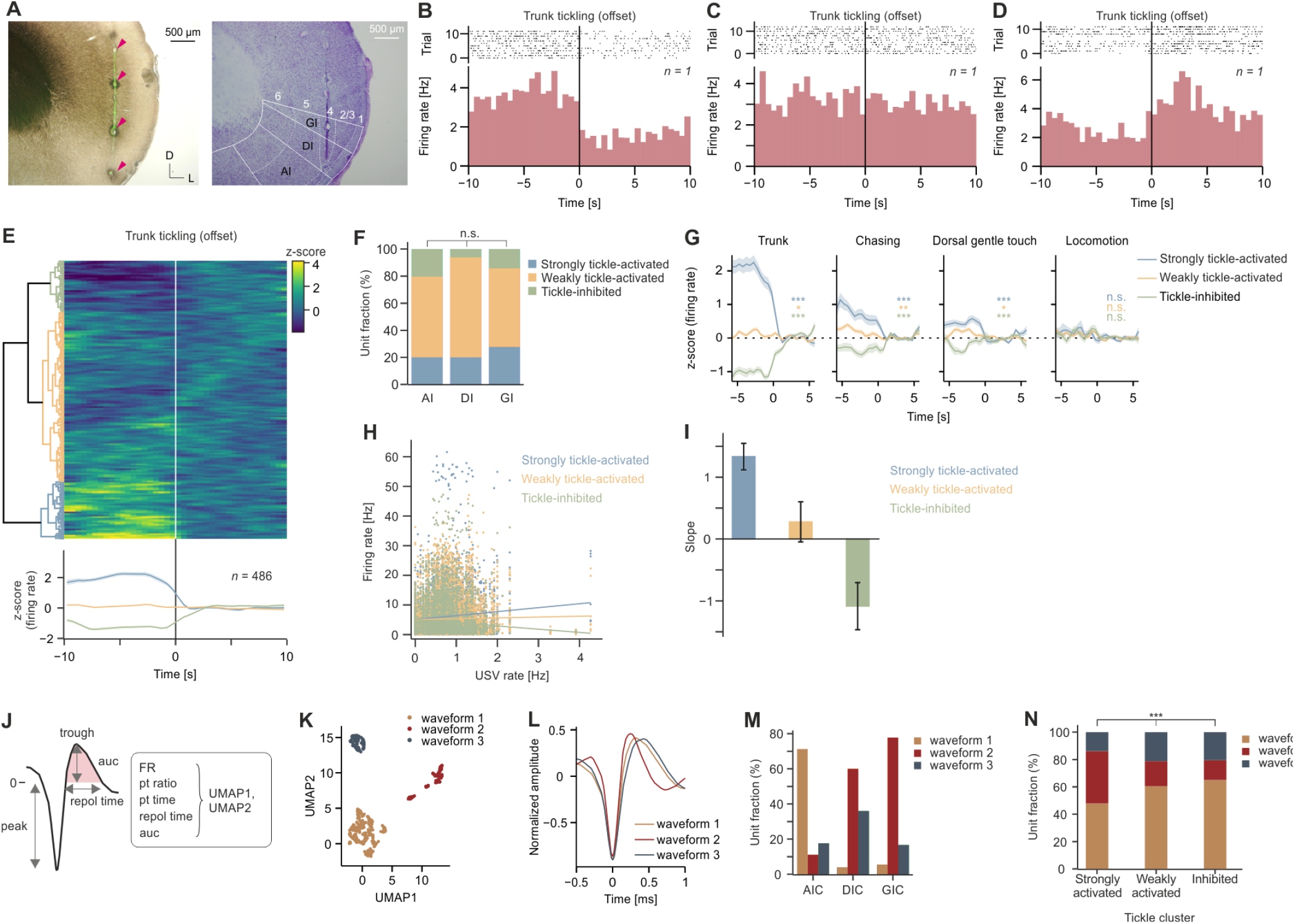
Differential representation of playfulness in aIC neurons. **A)** Identification of electrophysiological recording sites. Left: coronal section of anterior insular cortex at 2.76 mm anterior from bregma (green: tetrode track; arrowheads: lesions; D: dorsal; L: lateral). Right: same section stained with Nissl. Numbers indicate layers. GI: granular insula; DI: dysgranular insula; AI: agranular insula. **B-D)** Peristimulus raster plot (top) and time histogram (PSTH, bottom) of firing rate in three representative units, aligned to the offset of trunk tickling. Data are binned to 500 ms. *n* indicates number of units. **E)** Clustering of units based on PSTHs upon trunk tickling offset. Top, heatmap shows z-scored firing rate of each unit, sorted according to hierarchical clustering (dendrogram, left). Bottom, mean z-scored firing rates ± SEM of clusters. *n* indicates number of units. **F)** Anatomical distribution of each cluster across aIC subregions. *p* = 0.052 (Chi-square test of independence). **G)** PSTHs of firing rate of clusters from E) aligned to the offset of different events. Data obtained from 29 sessions, 7 animals. Statistical comparison was performed using one sample t-test of z-score in [-5, -1] s, with population mean of 0 as null hypothesis. **H)** USV rate vs. firing rate during break periods. Each data point represents a unit in a given break event. Data are fitted with linear mixed effect models for each cluster. **I)** Slope of linear regressions from H), mean ± SEM. **J)** Schematic average of normalized spike recorded extracellularly (left). Waveform features (right) were used for UMAP. FR: firing rate; pt: peak to trough; repol: repolarisation; auc: area under curve. **K)** Clustering of UMAP1 and UMAP2 of waveform features. Clustering was performed using KMeans. **L)** Average waveforms of waveform clusters. **M)** Anatomical distribution of each waveform cluster in K). **N)** Waveform distribution of tickle clusters in K). *p* < 0.001 (Chi-square test of independence).

#### Histology

After the last recording, animals were anaesthetized with ketamine (88 mg/kg body weight)/xylazine (7.5 mg/kg body weight) and the tetrode/probe tracks were labelled with electrolytic lesions by applying a DC current (8 s, 8 μA, electrode tip negative; using nanoZ, White Matter LLC, USA). Animals received an overdose of isoflurane and were transcardially perfused with a pre-fixative phosphate buffer solution followed by a 4% paraformaldehyde solution. The brains were dissected and post-fixed in 4% paraformaldehyde overnight. For histological processing, brains were cut in 100 μm coronal sections and Nissl stained for the assignment of recording sites to layers and observed under epifluorescent microscope (BZ-8000, Keyence, Japan). Lesions were also discernible in brightfield images (Figure 2A). Furthermore, to ensure accurate track positioning, we examined the DiI or DiO traces under the same epifluorescent microscope.

### Pharmacology

#### Cannula implantation

After two weeks of tickling habituation, the bilateral implantation of 26-gauge guide cannulae (C315GAS-5/SPC, Bilaney Consultants GmbH, Dusseldorf, Germany) into the insular cortex followed the procedure outlined in the “Drive and Implantation” section. The depth was consistently set at 3.5 mm from the pia level. The cannulae were securely fixed using dental cement (Paladur, Heraeus Kulzer, Hanau, Germany) capped with dummy cannulae to prevent any foreign objects from entering. Subsequently, rats were given a recovery period of one to two days.

#### Infusions

A 33-gauge internal locking cannula (C315LI/SPC, Bilaney Consultants GmbH, Dusseldorf, Germany) was inserted into the guide cannula with a protrusion of 1 mm making the infusion of solutions at 4.5 mm from pia level. Internal cannulae were connected to 5 µl Hamilton micro syringe by polyethylene (PE-20) tubing. Rats underwent bilateral infusion with a volume of 500 nl of either muscimol (400 ng/µl) (Sigma-Aldrich, M1523), gabazine (25 pmol) (Sigma-Aldrich, S106), or saline at a rate of 250 nl/min using a stereotaxic injector (Stoelting, USA). A period of 5 minutes was allowed for diffusion following infusion. Tickling experiments began one hour after muscimol infusion and 10 minutes after gabazine infusion. The experimenter responsible for handling the animals remained blind to the experimental conditions throughout the procedures. Following the final recording session, DiI was infused to confirm the opening of the cannulas and to assess the diffusion of the drugs, although drug diffusion dynamics may differ from those of DiI.

#### Histology

Animals received an overdose of isoflurane and were transcardially perfused with a pre-fixative phosphate buffer solution followed by a 4% paraformaldehyde solution. The brains were removed and post-fixed overnight. Slices (80 µm thick) were collected throughout the anterior insula and analysed under an epifluorescent microscope to precisely identify the locations of infusion sites and DiI diffusions. Only animals with bilateral tracks accurately terminating within the insula were included in the final analysis.

#### Analysis

We quantified the USV rate during trunk (dorsal and ventral) tickling for each session. The average USV rate for sessions under each drug condition (muscimol and gabazine) was normalized by divided it by the average USV rate from saline sessions for the same subject. One-sample t-test was used to evaluate the normalized USV rates, with the population mean set to 1 to assess deviations from the saline condition. For the analysis of hand chase behavior during no-contact play phases, the duration of hand chasing was measured and expressed as a percentage of the total no-contact play phase duration for each session. Statistical evaluation of these chasing duration percentages across drug conditions was conducted using a linear mixed effects model, with subject ID as a random intercept to account for inter-subject variability in the baseline chasing behavior. The rate of playful jumps during break periods was quantified for each session and subsequently evaluated using a linear mixed effects model to account for inter-subject variability.

### Anterograde tracing

#### Virus and injections

For anterograde tracing from the insula, two 5 weeks old rats habituated to tickling underwent unilateral injection of 100 nl at 30 nl/min of AAV2/5.hsyn.eGFP (Addgene, 50465-AAV5) in the insular cortex at the following coordinates: AP: + 2,76 mm; ML: 4,5 mm; Bregma DV: 5 mm. Surgery procedures were identical to those described earlier. Pipettes with a tip diameter of 22 µm were used for the injections. The pipettes were backfilled with the virus until one third of their volume, after which they were backfilled with oil. Following the injection, a diffusion period of 20 minutes was allowed before gradually retracting the pipette to minimize any potential overflow. Animals were allowed a recovery period of 4 weeks before being perfused to enable virus expression. During this time, animals were kept being tickled.

#### Histology and c-Fos staining

Four weeks after viral injection, and after the last tickling session by one hour, animals were transcardially perfused and the brains were collected. Sections of 80 µm thickness were obtained. Tissue was then washed with PBS 1X then blocked for nonspecific antibody binding by incubation for 2 hours at room temperature in blocking solution containing 1% Triton/10% normal goat serum (NGS) in PBS 1X. Sections were then incubated in a 0.1% Triton, 5% NGS solution containing primary rabbit anti-cFos antibody (1:1000; Cell Signalling, 2250) for 48 hours at 4°C. The secondary donkey anti-rabbit Alexa plus 555 antibody (1:500; Life Technologies, A-32794) along with anti-GFP DyLight 488 (1:500; BioTechne, NBP1-69969) was then applied at room temperature for 2 hours. Nuclei were stained with 4′, 6-diamidino-2-phenylindole (DAPI) (1:10; Carl Roth, 6335.1). Imaging was performed with an epifluorescent microscopy (VS200, Olympus, Japan). Intensity of axonal projections from the anterior insula as well as c-Fos expression in various brain areas was semi-quantitatively assessed.

### Statistics

Intergroup comparisons were performed with signed-rank test for paired data, and rank-sum test for unpaired data. *n* refers to sample size. Data were analysed using MATLAB 2021a and Python 3.7.9.

## Results

### The anterior insular cortex responds to tickling with symmetric firing patterns and diverse physiological properties

Rats emitted 50 kHz USVs during the tickling sessions (Figure 1C), in line with previous report [27]. During the intro periods, rats exhibited minimal USVs (0.02 [0.0, 0.07] Hz; median & IQR), albeit occasional occurrences, suggesting anticipatory effects from habituation, coupled with the rewarding nature of tickling [28]. Trunk (dorsal and ventral) tickling elicited the highest USV rate while gentle touch induces significantly lower USV rate (1.49 [0.89, 2.05] Hz; median & IQR) than trunk tickling (2.90 [2.66, 3.61] Hz; median & IQR; *p* < 0.001), indicating the scalability of emotions. Hand chasing during no-contact play phases also triggered USVs (2.74 [2.45, 3.65 Hz; median & IQR) despite the absence of direct tactile stimulation. These findings indicate that 50 kHz USVs can be triggered by diverse stimuli, highlighting the generalization of emotions. Additionally, rats emitted USVs upon playful jumps (i.e. *Freudensprung* [27]) during break periods (1.70 [1.36, 2.41] Hz; median & IQR). However, USV rate during locomotion, defined by the rat’s movement within the arena during the intro period, was negligible (0.00 [0.00, 0.00] Hz; median & IQR), suggesting that USVs emitted upon jumping were not merely a byproduct of motor movement i.e. the specificity of emotions. Furthermore, USV rates during break periods when the rat was alone in the arena were higher than the intro period (0.77 [0.53, 0.95] Hz; median & IQR; *p* < 0.001), unaffected by tactile or visual stimuli from the experimenter. The USV emission diminished towards the end of the session during the outro phase (0.20 [0.08, 0.36] Hz; median & IQR) but remained higher than in intro period (*p* < 0.001), implying the persistence of emotions. Thus, we confirmed that 50 kHz USVs serve as reliable indicators of playful emotions during tickling sessions.

To investigate the role of the anterior insula (aIC) in playful emotions, we recorded extracellular neural activity across layers 2/3 through 6 across subregions of the aIC (Figure 2A). Our findings revealed diverse activity patterns within the aIC: while some units demonstrated a strong increase in firing rate during trunk tickling, others displayed a decrease in activity, and a subset showed minimal discernible response (Figure 2B-D). These distinct activity profiles were categorized through hierarchical clustering of unit activity during trunk tickling (Figure 2E). Approximately 20% (100 out of 486 total units) units strongly increased activity during tickling, and a similar proportion showed inhibition (99 out of 486 units). Weakly-activated units constituted the majority, accounting for approximately 59% (289 units) of the total recorded units. The observed clusters did not exhibit any specific anatomical distribution within the aIC subregions (Figure 2F). To understand whether these activities represent tactile, locomotive, or playful emotional response, we tracked the firing patterns of the clusters, obtained based on their response to tickling, across other interaction phases (Figure 2G). Our analysis revealed that during hand chasing behavior and dorsal gentle touch, the activity patterns of the clusters remained consistent. On the other hand, none of the clusters responded to locomotion during the intro period. Thus, similar to USVs, the aIC activity observed during tickling extended beyond mere tactile-based activity or locomotion; instead, it appears to be specific to the context of playfulness and generalized across playful interactions.

Given our observations indicating similar modulation patterns of USVs and neuronal activity in the aIC during playful interactions, we sought to investigate a potential causal link between USVs and the aIC activity. Accordingly, we examined the relationships between USV rates and firing rates of aIC neurons for each tickle-response cluster (Figure 2H-I). To eliminate potential confounding effects of aIC neuron coactivation and USVs induced by touch, we specifically restricted this analysis to break periods. Using linear mixed effects models and accounting for individual unit baseline firing rates as random effects, we revealed a positive correlation between the firing rate of the tickle-activated cluster units and USV rate. Specifically, for the strongly tickle-activated cluster units, we observed that an increase in USV rate by 1 Hz corresponded to an increase in firing rate of the units by 1.33 Hz. Conversely, the weakly tickle-activated cluster units displayed a weak correlation with USVs. Furthermore, the tickle-inhibited cluster units exhibited a negative correlation with USVs: a 1 Hz increase in USV rate corresponded to a decrease in firing rate by 1.09 Hz. These findings, underscored by the scalable relationship between firing rates and USV rates, collectively suggest a differential involvement of aIC neuronal clusters in dynamically modulating the intensity of playful emotions.

The diverse activity patterns observed in the aIC hinted at the presence of distinct neuronal populations. To test this notion, we performed waveform analysis to investigate whether the recorded units exhibited heterogeneity or homogeneity. We extracted spike waveform features, such as firing rate, peak-to-trough ratio, and others, from all recorded units (Figure 2J). Subsequently, we performed dimensionality reduction by projecting these features into UMAP space (Figure 2K). Clustering using K-Means algorithm unveiled three distinct waveforms with varying proportions across different anatomical locations within the insula (Figure 2L-M). Furthermore, we investigated how these waveforms contributed to the formation of the tickle response clusters. Our analysis revealed differences in waveform composition among the three tickle response clusters (Figure 2N). For instance, the cluster strongly activated by tickling exhibited a higher fraction of waveform 2 compared to the tickle-inhibited cluster.

### Anterior insular global pharmacological manipulation yielded no discernible effects on play behavior

Based on the identified activity patterns within the aIC in response to playful contexts, we aimed to understand its precise contributions to the observed behavior. Therefore, we bilaterally implanted cannulas in the aIC for comprehensive manipulation with drug infusion (Figure 3A). Through infusions of muscimol, a GABA-A receptor agonist, and gabazine, a GABA-A receptor antagonist, we achieved aIC inhibition and disinhibition, respectively. Interestingly, neither inhibition nor disinhibition had any discernible effect on play behavior in rats during tickling, as assessed by normalized USV rate to saline infusion sessions, chasing duration during no-contact play phases, and jump rate during break phases (Figure 3B-D).

**Figure 3:**
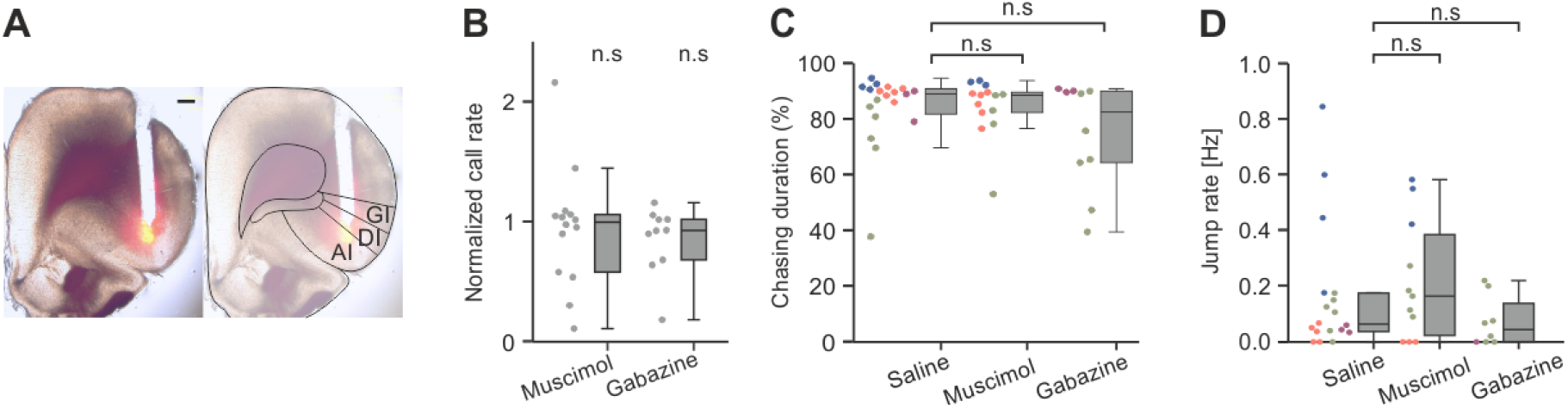
Pharmacological manipulation of aIC. **A)** Coronal section of cannula implantation site in aIC. Red fluorescence represents DiI injection before perfusion. AP: 2.76. Scale bar = 0.5 mm. **B)** Ultrasonic vocalization rate during tickling sessions normalized to mean call rate in saline controls for each subject (muscimol: *p* = 0.667, *n* = 14 sessions; gabazine: *p* = 0.126, *n* = 10 sessions; one sample t-test). **C)** Chasing duration during no contact play (muscimol vs. saline: *p* = 0.75, *n* = 14 sessions; gabazine vs. saline *p* = 0.33, *n* = 10 sessions; linear mixed effects model). Colors indicate subjects. **D)** Jump rate after trunk tickling (muscimol vs. saline: *p* = 0.62, *n* = 11 sessions; gabazine vs. saline *p* = 0.43, *n* = 8 sessions; linear mixed effects model). Colors indicate subjects.

### The anterior insula projected to areas activated by tickling

Given the lack of observed effects on play behavior following global pharmacological manipulation, our focus shifted towards delineating specific circuits of the aIC. Thus, we performed anterograde tracing from the aIC, where we observed play representations, to identify its downstream projection target areas. Our results showed a rich network of projections with diverse densities, extending to numerous brain regions on both ipsilateral and contralateral sides (Table 1). The integration of tracing with post-tickling c-Fos staining revealed that these aIC projections targeted several brain regions associated with tickling-induced activation, such as the nucleus accumbens, the mediodorsal thalamic nuclei, the posterior thalamic nucleus, and the amygdala (Figure 4).

**Table 1:**
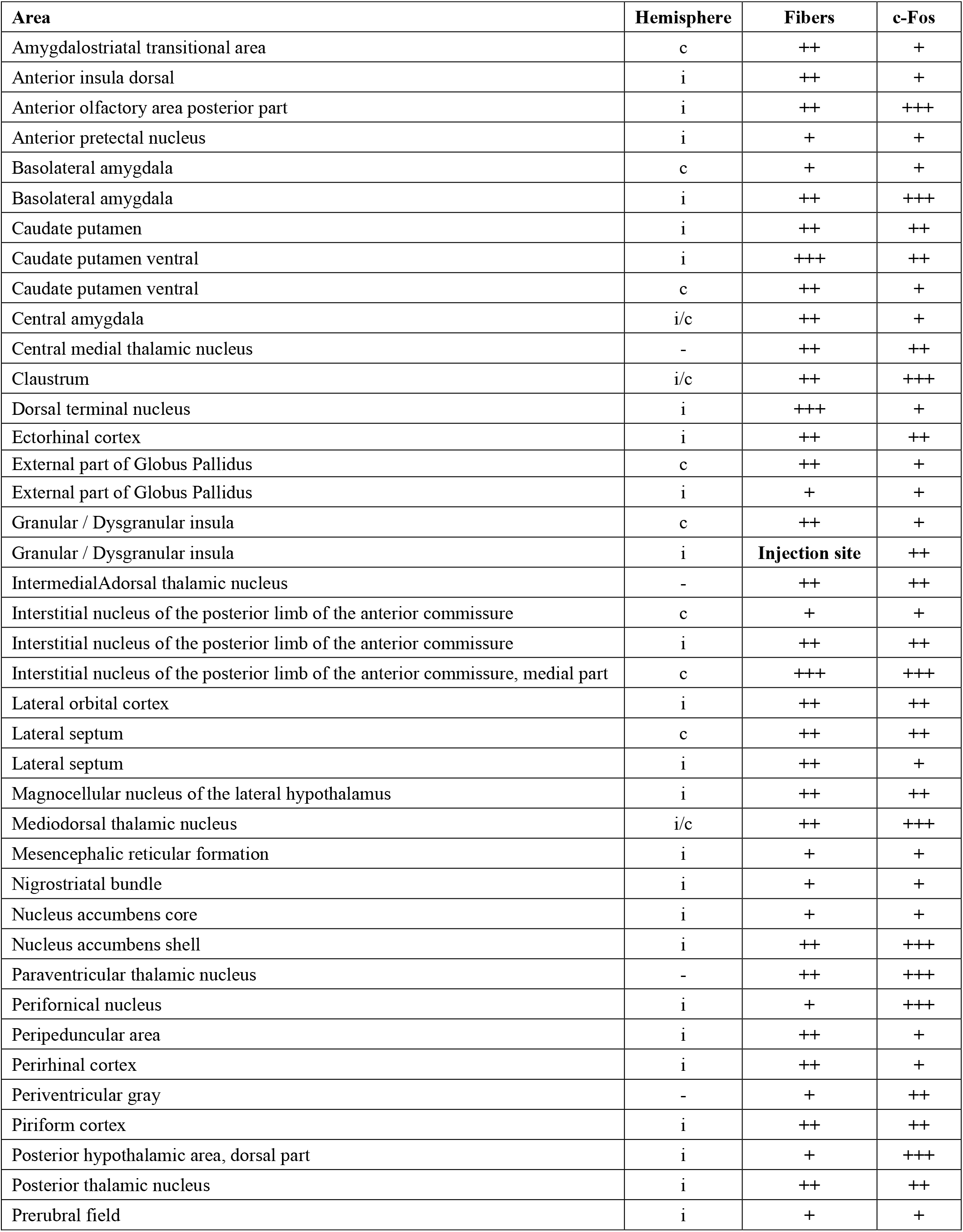
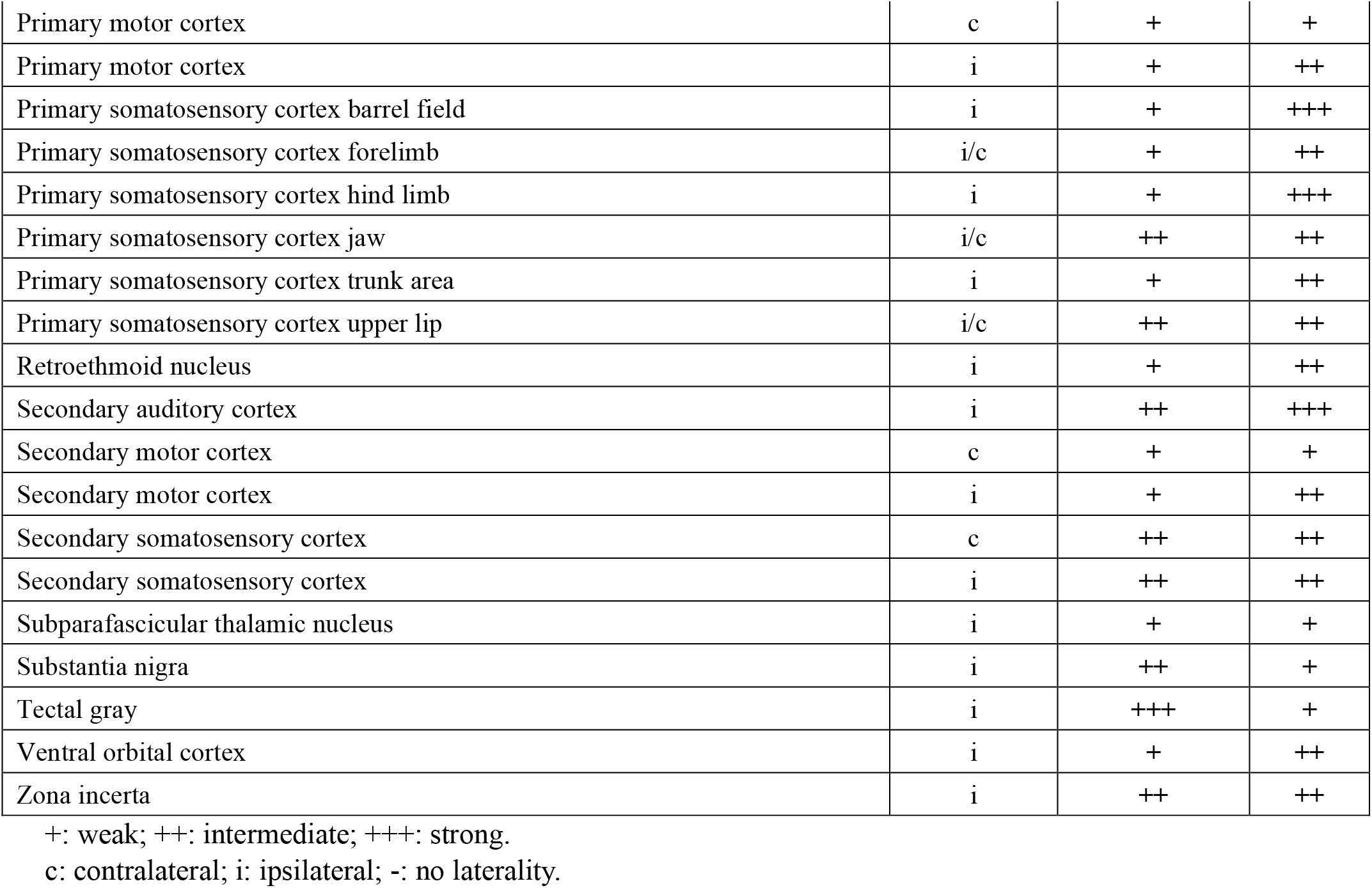
Anterograde tracing from the aIC and c-Fos after tickling colocalization.

**Figure 4:**
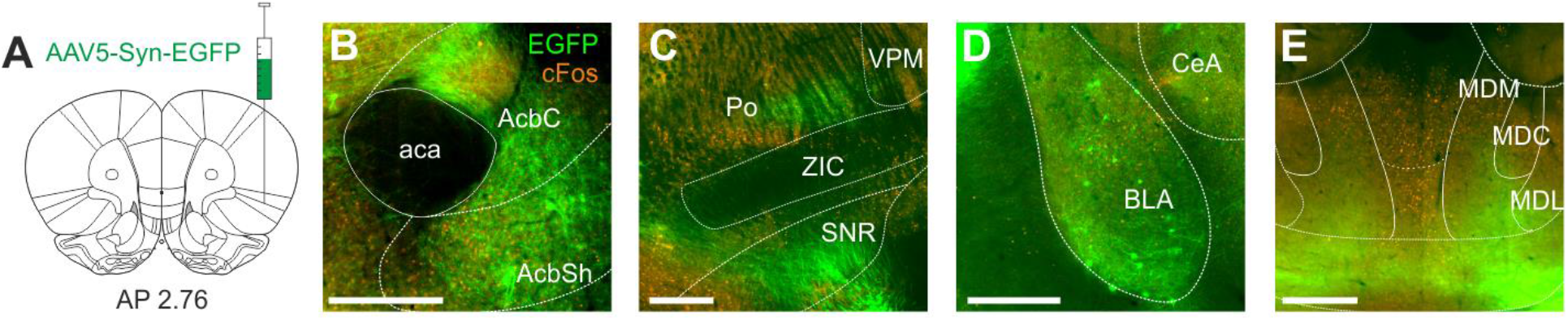
Anterograde tracing from aIC and c-Fos colocalization. **A)** Anterograde viral injection in aIC using AAV5-Syn-EGFP. **B)** GFP labelled fibers and cFos expression. aca: anterior commissure, AcbC: accumbens core, AcbSh: accumbens shell. AP: 3.00, ipsilateral. **C)** Same as B); Po: posterior thalamic nucleus, VPM: ventral posteromedial nucleus, ZIC: zona incerta caudal, SNR: substantia nigra reticular. AP: -4.56, ipsilateral. **D)** Same as B); BLA: basolateral amygdala, CeA: central amygdala; AP: -1.76, ipsilateral. **E)** Same as B); MDM: mediodorsal thalamic nucleus medial, MDC: mediodorsal thalamic nucleus central, MDL: mediodorsal thalamic nucleus lateral. AP: -3,36, ipsilateral and contralateral. Scale bars: 0.5mm.

## Discussion

In this study, we aimed to dissect the role of the rat anterior insula in the processing of playful emotions, particularly those evoked by tickling interactions with humans. We uncovered a spectrum of activity patterns, physiological diversity, and projection targets within the aIC. Surprisingly, global manipulation of the aIC failed to produce observable effects on play behavior, prompting a deeper exploration into more specific insular circuits. The integration of anterograde tracing with post-tickling c-Fos staining suggests the involvement of diverse downstream regions, including the nucleus accumbens, the mediodorsal thalamic nuclei, and the amygdala, in mediating tickling-induced emotional responses.

The exploration of internal states or emotions is currently a prominent frontier in neuroscience and has been marked with remarkable advances. Investigations into aggressive states have spotlighted key neural substrates including the medial amygdala, bed nucleus of the stria terminalis (BNST), and ventromedial hypothalamus [53, 54]. Moreover, the study of pleasure-related phenomena has directed attention towards the nucleus accumbens, anterior cingulate cortex, ventral pallidum, and amygdala [55]. The insular cortex, on the other hand, has recently emerged as a key structure for a spectrum of behaviors and affective processes, modulating feeding behavior [56-59], motivational processes [60], nociceptive processing [61, 62], saliency attribution [63, 64], anxiety [38, 65, 66], and social approach behaviors [67-70]. However, a major challenge in the field lies in objectively quantifying emotions in non-verbal species or animals, where quantification of emotions can be highly subjective. To overcome this, the identification of emotions relies on features such as valence, generalization, specificity, persistence, and scalability as previously described [30]. These emotional characteristics are studied by measuring the changes that they induce for instance in locomotive activity, vocalizations, endocrine responses, or facial expressions [30, 32, 41]. In our study, we utilized ultrasonic vocalizations (USVs) as indicators of internal states in tickled rats, fulfilling the features of emotions. USVs at 50 kHz are associated with positive valence [71-73] and, in our tickling sessions, were emitted across various playful contexts, including tickling, chasing, jumping, and gentle touch (Figure 1C). Moreover, USV emission was scalable between these contexts and specific to the emotional state induced by the playful interactions. In fact, previous studies have also demonstrated that these USVs are not emitted during tickling under anxiogenic conditions [10, 27], further supporting their role as indicators of positive internal states. Our analysis also revealed representations of emotional persistency in USVs. Thus, we demonstrate that 50 kHz USVs reliably signal playful emotions in response to tickling, aligning with key emotional attributes.

The response patterns observed in both tickle-activated and inhibited units within the aIC are indicative of a broader mechanism underlying the processing of playful emotions (Figure 2). Despite their distinct symmetric neurophysiological effects—activation and inhibition—these units collectively demonstrated generalizability and specificity in their response to playful contexts (Figure 2G). Moreover, the correlation of cluster activities with the emission of USVs during break periods (Figure 2H-I), albeit with opposite effects, suggests a complex interplay between these neuronal populations within the aIC in mediating and regulating the processing of fun and play emotions. The synchronized changes observed in both neuronal activity and behavioral responses, particularly in the linear modulation of USVs (Figure 2H-I), highlight a consistent relationship between brain function and observable behavior. This consistency emphasizes the neural foundation that mirrors the behavioral expressions indicative of internal states or emotions.

We revealed that distinct spike waveforms were associated functionally with tickle-response patterns and anatomically with substructures within the aIC (Figure 2L-N), suggesting that physiological and cellular underpinnings may account for the varied tickle response patterns. However, it is important to note that definitive confirmation of specific cell types exceeds the scope of our study. Without the utilization of proper molecular markers, any conclusions regarding cell types remain speculative.

Studies previously showed the strong activation of the trunk somatosensory cortex during tickling-induced vocalizations [27]. This aligns with our observation of increased firing rates in certain aIC units during tickling, suggesting a potential functional link between the somatosensory cortex and the aIC in mediating the emotional responses elicited by tickling. One possible interpretation is that the somatosensory cortex processes tactile and anticipatory sensations associated with tickling, while the aIC integrates this sensory input with emotional and motivational signals. This mechanism of associating subjective emotions to tactile sensations has also been demonstrated in pain processing in the insula [44, 62]. We previously demonstrated that activation in the deep-layer trunk somatosensory cortex evokes 50 kHz USVs [27], and tickling-induced ambivalence is represented in layer 5 but not layer 4 trunk somatosensory cortex [28]. Our anterograde tracing revealed the aIC projection to the posterior thalamic nucleus (Figure 4C), which is known to project to the layer 5 primary somatosensory cortex, mediating decision-making processing [74]. Hence, it could be speculated that the aIC neurons might mediate the deep-layer trunk somatosensory cortex via the posterior thalamic nucleus. Thus, a possible reciprocal loop between the somatosensory cortex and the aIC might underlie the intricate processing of sensory inputs and playful emotional responses. Another recent study showed the periaqueductal gray (PAG) revealed diverse and strong modulation of PAG units during tickling, play, and USV emission [10]. Although our study did not assess direct connectivity between the primary somatosensory cortex, the aIC, and PAG, it is still reasonable to speculate that interaction among these regions might be characterized by feedback loops and parallel processing pathways contributing to the observed behavioral output in terms of USVs and playfulness.

The observation that pharmacological manipulation of the aIC did not directly impact playfulness (Figure 3), unlike the blockade of other brain regions like the PAG [10], nucleus accumbens [75], and habenula [76], suggests that the aIC may not be necessary as a whole to mediate playfulness. The insula is known for its extensive connections with various brain regions involved in sensory, emotional, and cognitive processing, suggesting its role in integrating diverse inputs to modulate behavior [43]. This complexity is further underscored by the presence of distinct response patterns during playful contexts within the aIC, which may regulate playfulness in opposing ways. Manipulating the unspecific neurons in the aIC may not yield a specific effect on playfulness due to the potential balance of excitatory and inhibitory influences from different neurons. The concept of functional specificity is evident in other brain regions, such as the hypothalamus, where adjacent cells can mediate vastly different behaviors like aggression, mating, and parenting [77, 78]. Similarly, within the insula, the localization of neurons in different layers can result in opposing effects on behaviors like appetitive drinking, depending on their projection targets. For example, layer 5 neurons in the insula can mediate both promotion and inhibition of drinking behavior, with layer 5a and 5b neurons projecting to different downstream targets with distinct functional outcomes [79]. These results may imply that specific pathways or downstream areas within the insula may play a more crucial role in regulating behaviors, suggesting that the distinct tickle-response clusters may have distinct projection targets. Therefore, it would be crucial to identify and selectively target these specific pathways that differentially mediate playfulness.

The anterograde tracing results, in line with other studies [80], revealed extensive projections from the aIC to various brain regions, including the mediodorsal thalamic nucleus, amygdala, and nucleus accumbens (Figure 4 & Table 1). These brain regions, known for their roles in social behavior, emotional regulation, and reward processing, have been extensively studied in the context of various behaviors, including play [8, 66, 81-87]. The interactions between these regions may contribute to the multifaceted nature of play behavior, including sensory enjoyment, emotional engagement, and motivational drive. Therefore, further studies are needed to dissect the functional contributions of individual pathways within this circuit.

In conclusion, through a combination of electrophysiological recordings, neuroanatomical tracing, and behavioral assays, we uncovered a spectrum of physiological responses and neuronal diversity within the anterior insular cortex involved in the processing of playful emotions evoked by tickling in rats. Our results offer a high-resolution glimpse into cellular-level processes contributing to the complex interplay between brain function and behavior in non-verbal species in naturalistic behaviors.

## Acknowledgments

Authors thank Jannik Wagner for assisting software development, Richard Necel, Sascha Amenta, Stefan Schön and Michael Marx for technical support, Ryu Gyo for technical advice, and Alan Kania for microscope assistance. Authors also thank Tobias Ruff, Hiroaki Norimoto, Yan Tang, and Juan Ignacio Sanguinetti-Scheck for discussions and comments.

## Author contributions

Conceptualization, S.D. and S.I.; Experiments, S.D. and S.I., Formal analysis, S.D. and S.I.; Visualization, S.D.; Funding acquisition, S.I.; Supervision, S.I.; Writing-original draft, S.D., and S.I.

